# *Azospirillum brasilense* and cytidine as combination treatment to enhance drought resilience of peas

**DOI:** 10.64898/2026.10.26.761419

**Authors:** Bidya Ojha, Janice Tagoe, Shelley M. Horne, Birgit M. Prüß

## Abstract

**Aims:** This study tested the efficacy of two combination treatments on drought resilience of peas; *Azospirillum brasilense* Sp7 with the chemoattractant seedling exudate compound cytidine or pea seedling exudate.

**Methods and Results:** Phenotypic characterization revealed that drought tolerance of peas treated with ACyt was 68.9% higher than that of peas treated with the inoculant alone (A). Dry shoot weight of ACyt Drought plants was 30.7% higher than that of plants from the untreated control group NB Drought, drought recovery of ACyt plants was 21.6% better. The treatment with seedling exudate (AEx) was less effective.

16S rDNA analysis revealed that increase in drought tolerance was accompanied by modulation of the rhizosphere. The abundance of *Azospirillum* sp. was 954% higher in ACyt Drought plants than in NB Drought ones. The ACyt treatment may contribute to protecting pea plants from drought by increasing the abundance of *Azospirillum*. Within the complete analysis of the 16S rDNA data set, both ACyt Drought and AEx Drought groups had similar increases in Shannon and Pielou alpha-diversity indices, relative to the NB Drought group. These two test groups also exhibited similar changes in the bacterial community structure.

**Conclusion:** The ACyt treatment enhanced drought tolerance and recovery. Both treatments enhanced alpha-diversity and modulated the rhizosphere.

**Impact statement:** Drought is a major constraint on pea production worldwide, necessitating sustainable approaches to improve crop resilience. This study demonstrates that the combined application of *A. brasilense* Sp7 and cytidine (ACyt) and to a lesser extent that of the inoculant with seedling exudate (AEx) enhance drought tolerance and recovery in peas. Both treatments significantly alter rhizosphere bacterial community composition. The study identifies microbial community shifts that accompany enhanced drought resilience and provides a foundation for future studies investigating the mechanisms underlying plant-microbe interactions and stress tolerance in peas.

## Introduction

Field pea (*Pisum sativum* L.) is a globally cultivated member of the family Fabaceae, legumes. A unique feature is the ability to form a symbiosis with root nodulating *Rhizobium leguminosarum* bv. *viciae,* in which the bacterial contribution is the fixation of atmospheric nitrogen (Lindström and Mousavi 2020). The full productive potential of peas is increasingly compromised by an array of environmental pressures; drought stress alone has been estimated to suppress pea yields by as much as 40% under severe conditions (Sorensen et al. 2002).

Considerable effort has focused on improving drought tolerance of plants. This includes plant breeding, inoculation with plant growth-promoting rhizobacteria (PGPR), and diverse modulations of the biochemical and molecular make-up of the plants (Haghpanah et al. 2024). While less investigated, post-drought recovery has increasingly been recognized as a key determinant of plant performance and resilience (Zheng et al. 2023; Wang and Xu 2026).

*Azospirillum brasilense* is a well characterized PGPR. It exerts its beneficial effects through enhancement of root architecture, improved acquisition of water and nutrients, increased phytohormone biosynthesis, emission of volatile compounds, nitric oxide signaling, and the priming of systemic defense pathways (Creus et al. 2005; Cassán 2026). The deployment of *A. brasilense* specifically within pea cropping systems, particularly in the face of environmental stress scenarios, has received limited experimental attention (Ejaz et al. 2020; Fourneau et al. 2024; Nisha et al. 2024). For this study, we use *A. brasilense* Sp7 as the first component of the combination treatment.

The second component is cytidine, a nucleobase and seedling exudate compound (Nisha et al. 2024). Plant exudates can serve as chemoattractant signals that selectively attract and support specific microbial taxa that are beneficial to the plant (Gaworzewska and Carlile 1982; Armitage et al. 1988; O’Neal et al. 2020; Compton and Scharf 2021; Fourneau et al. 2024; Wang et al. 2026).

Recent investigations from our laboratory established that exudates from pea, tomato (*Solanum lycopersicum*), and cucumber (*Cucumis sativus*) seedlings contained nucleosides (Nisha et al., 2024). The ability of *A. brasilense* Sp7 to migrate towards cytidine was confirmed on semi-solid agar plates (Nisha et al. 2024). In a consecutive study, cytidine approximately doubled lateral root numbers (Nisha et al. 2025). It was decided to use *A. brasilense* Sp7 together with cytidine to improve resistance of tomatoes towards infection with *P. syringae* (Tagoe et al. 2026).

This study constitutes the second example of using *A. brasilense* Sp7 and cytidine to aid plants in resisting stress, this time against drought. As comparison, we included a second treatment, consisting of *A. brasilense* Sp7 and total seedling exudate. We measured drought phenotypes of pea plants for several days after withholding water at week 4 in a drought tolerance experiment. In a drought recovery experiment, we resumed watering on the 6^th^ day after withholding water. DNA was isolated from roots from one of the tolerance experiments and the 16S rDNA of bacteria was sequenced. Major modulations of the rhizosphere were observed in response to the treatments.

## Materials and Methods

### Drought experiment

The bacterial strain used in this study was *Azospirillum brasilense* Sp7 (ATCC 29145; DSMZ 1690; (Tarrand et al. 1978)), provided by Dr. Gladys Alexandre (University of Tennessee, Knoxville, TN). Plant experiments were done with CDC Striker peas (Warkentin 2004). Protocols for bacterial and plant growth are published at dx.doi.org/10.17504/protocols.io.6nvw6pn5wg mk/v1). Pea plants were grown at the Jack Dalrymple Agricultural Research Complex (https://www.ag.ndsu.edu/greenhouse) for 4 weeks. The drought study was conducted as two independent experiments on drought tolerance and drought recovery. For both experiments, pea plants were grown under watering for four weeks in six treatment groups: combinations of presence or absence of *A. brasilense* Sp7(A), cytidine (Cyt) or seedling exudate (Ex), and an untreated control (NB).

In the tolerance experiment, each of the six original treatment groups was split into drought and watered sub-groups, resulting in twelve groups of nine replicate plants. The recovery experiment used six treatment groups with nine replicate plants. At the beginning of week five, water was withheld from drought groups, while watered groups continued to receive water. During the drought period, plants were monitored daily for drought stress symptoms, including yellowing, leaf curling, and leaf shrinkage. Starting on day four of the drought period, plants were visually scored as percent health score. One healthy watered plant was compared with the drought stressed plant to give the scores. Examples of plants with shoot health scores in the range of 30-100% are included at dx.doi.org/10.17504/protocols.io.6nvw6pn5wgmk/v1. Scoring continued until day nine, when severesymptoms and plant decline were evident. On day 9, roots and shoots were collected in separate paper bags for dry weight determinations. Weights were expressed in g. For the drought recovery experiment, water was withheld from all nine replicate plants per group starting after four weeks. Plants were subjected to six days of drought, watering was initiated on day 6 with 50 ml of dH_2_O per plant every second day. Shoot health was visually scored from the onset of drought stress through the recovery period.

Data are presented as Box and Whiskers generated with GraphPad Prism 11.0.2 (92). Two-way ANOVA was used for statistical analysis of the shoot health scores, one-way ANOVA for the root/shoot weights. Tukey was used for pairwise comparisons. A *p*-value below 0.05 was indicative of a statistically significant difference between groups. Percentage increases were calculated using the formula: percent increase = ((mean of treatment group – mean of control group)/mean of control group) * 100.

### 16S rDNA analysis of the rhizosphere from the drought tolerance experiment

To study the bacterial community of the pea rhizosphere in response to the treatments, as well as under drought conditions, we analyzed 16S rDNA abundances on roots from four plants from one of the drought tolerance experiments, following the protocol at dx.doi.org/10.17504/ protocols.io.6nvw6pn5wgmk/v1. Sequencing was done at the Dr. Thomas Glass Biotech Innovation Core at NDSU (https://www.ndsu.edu/biotech-innovation-core). The resulting OTU abundance table was exported as Excel file, still containing the amplicon sequence variants (ASV). This table was filtered (Nikodemova et al. 2023) by using an inclusion cutoff of >500 reads across all treatment groups and replicates. Analyses included the abundances of *Azospirillum*, three indeces of alpha-diversity, and composition of the microbiome across test groups (dx.doi.org/10.17504/protocols.io.6nvw6pn5wgmk/v1). Data analysis was done on R-Studio 2025.05.0+496 (https:github.com/Ojha-B/Master-s-Thesis-Pea-Drought-Microbiome/tree/main). Pairwise comparisons are included in Figures and text when the *p*-value was <0.05.

## Results and Discussion

### Drought tolerance and recovery of pea plants

The objective of the drought tolerance experiment was to evaluate the effect of treatments on shoot health of pea plants under several days of drought stress. As comparison, we tested the combination of *A. brasilense* Sp7 with total pea seedling exudate. Drought stress was imposed upon four weeks old pea plants for another 9 days of plant growth.

Watered treatment groups showed very little decline in the shoot health score over the entire time of the experiment, scores stayed between 80 and 100% (Supplemental Fig. 1). For drought stressed groups, shoot health declined progressively between days 4 and 9 after watering was stopped (Fig. 1A). The exception was the ACyt Drought group, where scores decreased from day 4 to day 7 and then remained stable. On day 9, the shoot health score of plants from ACyt Drought was 68.9% higher than that from A Drought at a *p*-value <0.0001. On day 8, ACyt Drought yielded a 28.5% higher shoot health score with a *p*-value of 0.0004. Relative to NB Drought, shoot health scores from ACyt Drought were 35.8% higher at a *p*-value of 0.0161 on day 9.

**Figure 1.**
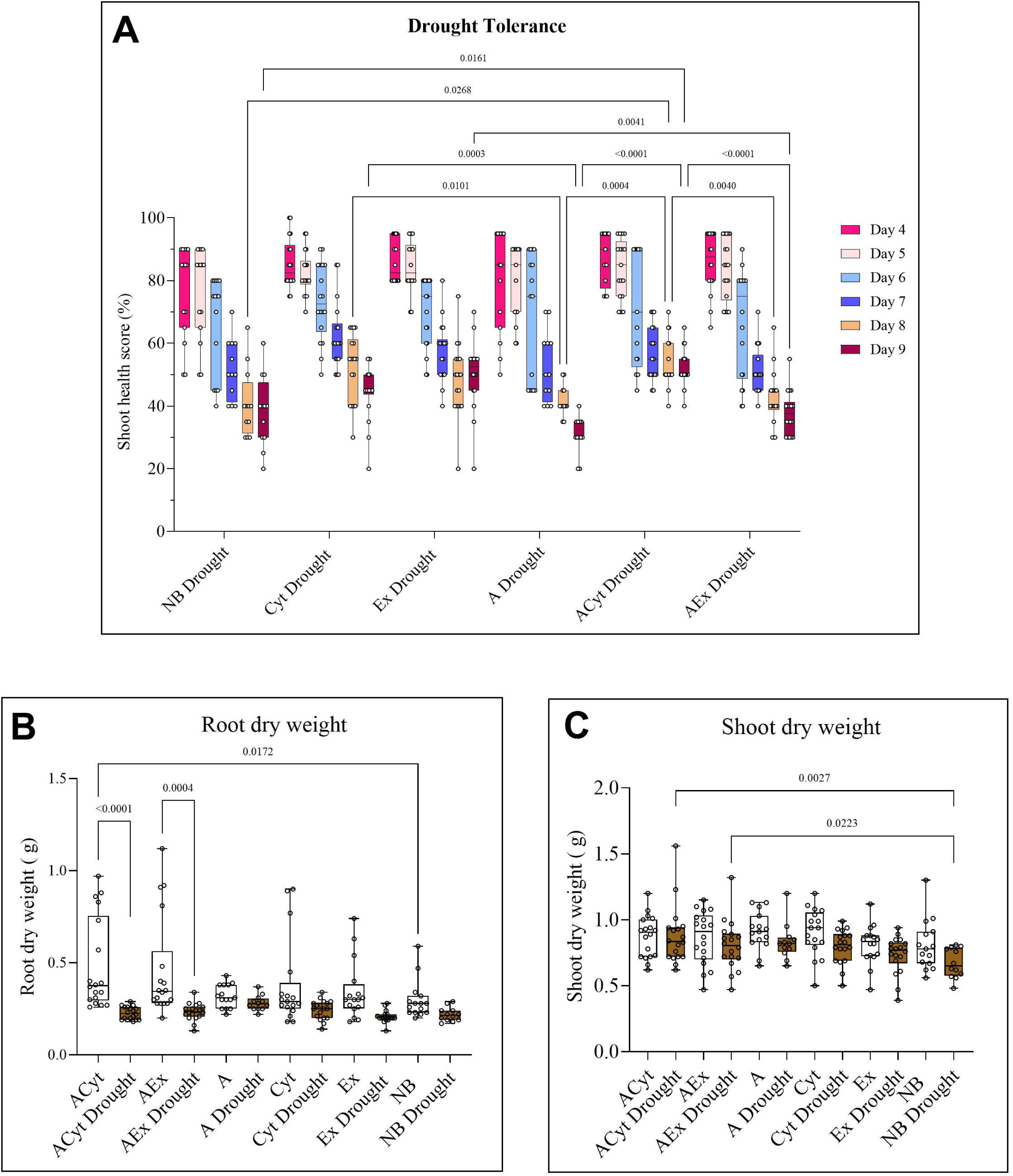
Drought tolerance experiment. Data were collected as percentage of healthy shoots from two replicate experiments over the drought period of days 4 to 9 (Panel A) and dry weights of roots (Panel B) and shoots (Panel C) from day 9. Treatment groups represent the presence or absence of *A. brasilense* Sp7 (A) and seedling exudate (Ex) or cytidine (Cyt) under drought conditions. NB is the untreated control. Lines connecting two bars indicate *p*-values from Tukey that are <0.05.

To correlate shoot health to dry mass of pea roots and shoots, we separated roots from shoots at day 9 post droughting and weighed them after drying (Figs. 1B and C). Dry weight of roots was larger in all comparisons between watered and droughted plants, in the cases of ACyt vs ACyt Drought and AEx and AEx Drought, this difference was of statistical significance (Fig. 1B). Among the drought treated plants, treatment did not significantly alter root dry weight. Shoot dry weights were significantly larger for plants from the ACyt Drought and AEx Drought treatment groups relative to NB Drought at 30.7% and 19.9%, respectively (Fig. 1C).

The rationale for the drought recovery experiment was to test the hypothesis that the more drought tolerant plants from the ACyt Drought and AEx Drought groups may also recover better. The drought tolerance experiment had shown that shoot health scores started to go into decline on day 6 without major differences between the treatment groups (Fig. 1A). It was decided to use day 6 to resume watering the pea plants for the drought recovery experiment.

As expected, shoot health scores during days 4 to 7 (Fig. 2) looked very similar to those from the drought tolerance experiment (Fig. 1). For all treatment groups, shoot health scores did not further decline after watering resumed, but stayed at least steady. Minor increases in shoot health scores upon re-watering were observed for treatment groups Cyt, A, and AEx. ACyt that had performed better under drought tolerance conditions was also superior at drought recovery. These plants were almost 100% healthy at day 10. From the statistical analysis of the day 10 data, ACyt exhibited a 21.6 % larger shoot health score relative to NB at a *p*-value of 0.0003. AEx shoot health relative to NB was larger by a mere 8.8% at a *p*-value that did not indicate statistical significance. At the phenotypic level, ACyt groups performed better than AEx groups.

**Figure 2.**
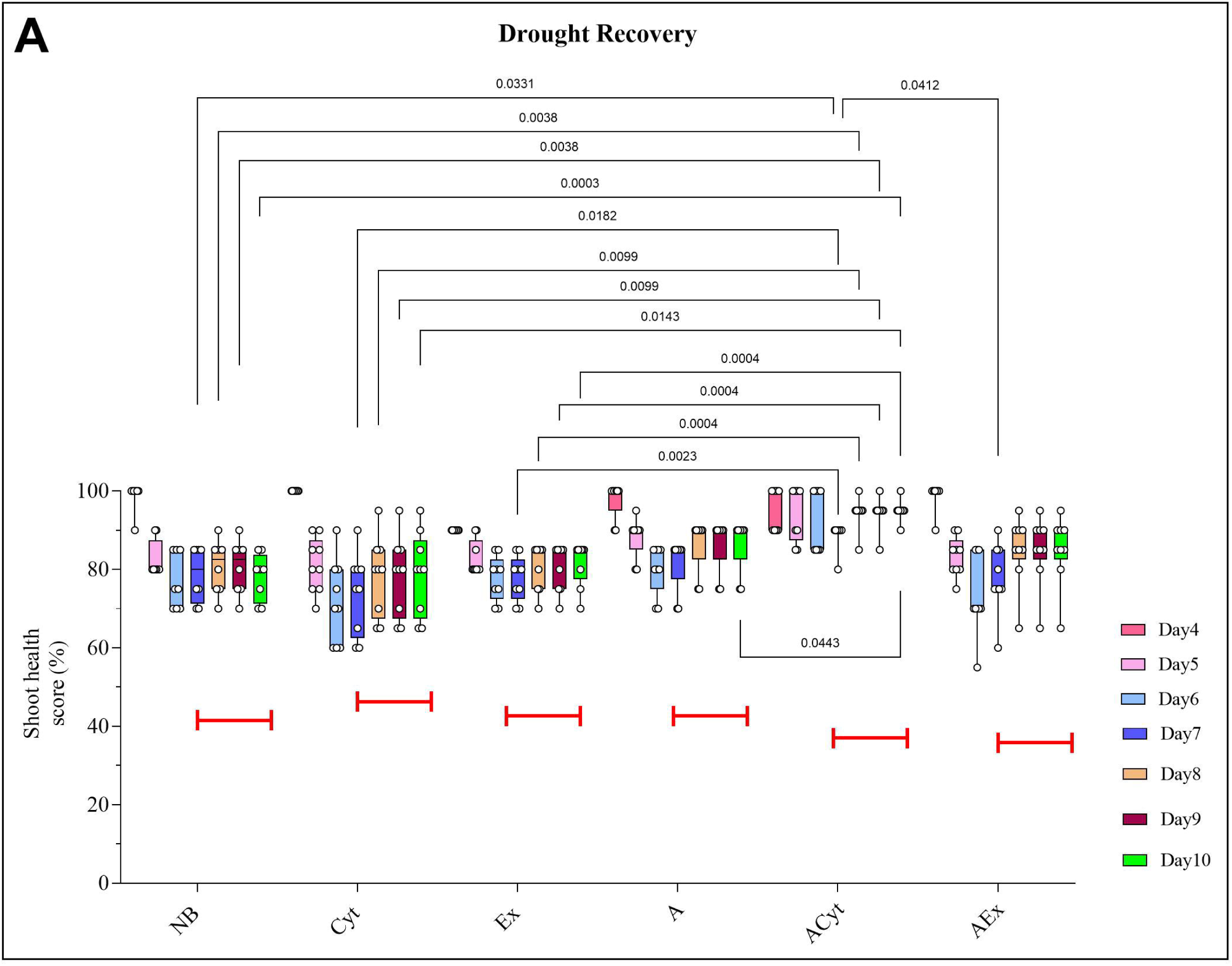
Drought recovery experiment. Shoot health scores were collected as percentage healthy shoots from one experiment across both the drought (days 4-6) and recovery (days 7-10) phases. Treatment groups represent the presence or absence of *A. brasilense* Sp7 (A) and seedling exudate (Ex) or cytidine (Cyt) under drought conditions. NB is the untreated control. Lines connecting two bars indicate *p*-values from Tukey that are <0.05.

To test the possibility that the superior performance of the ACyt treated may go in parallel with an increase in the abundance of the inoculant *A. brasilense* and other species of the *Azospirillum* genus, we used 16S rDNA sequencing from a rhizospheric homogenate. As expected, treatment groups inoculated with *A. brasilense* Sp7 showed detectable abundances of the inoculant, whereas un-inoculated treatment groups exhibited negligible abundances (Fig. 3A). The inoculant had an abundance that was 163% larger in ACyt Drought than in ACyt at a *p*-value of 0.0208. Abundance of *Azospirillum* sp., was 954% larger in ACyt Drought than in NB Drought (Fig. 3B).

**Figure 3.**
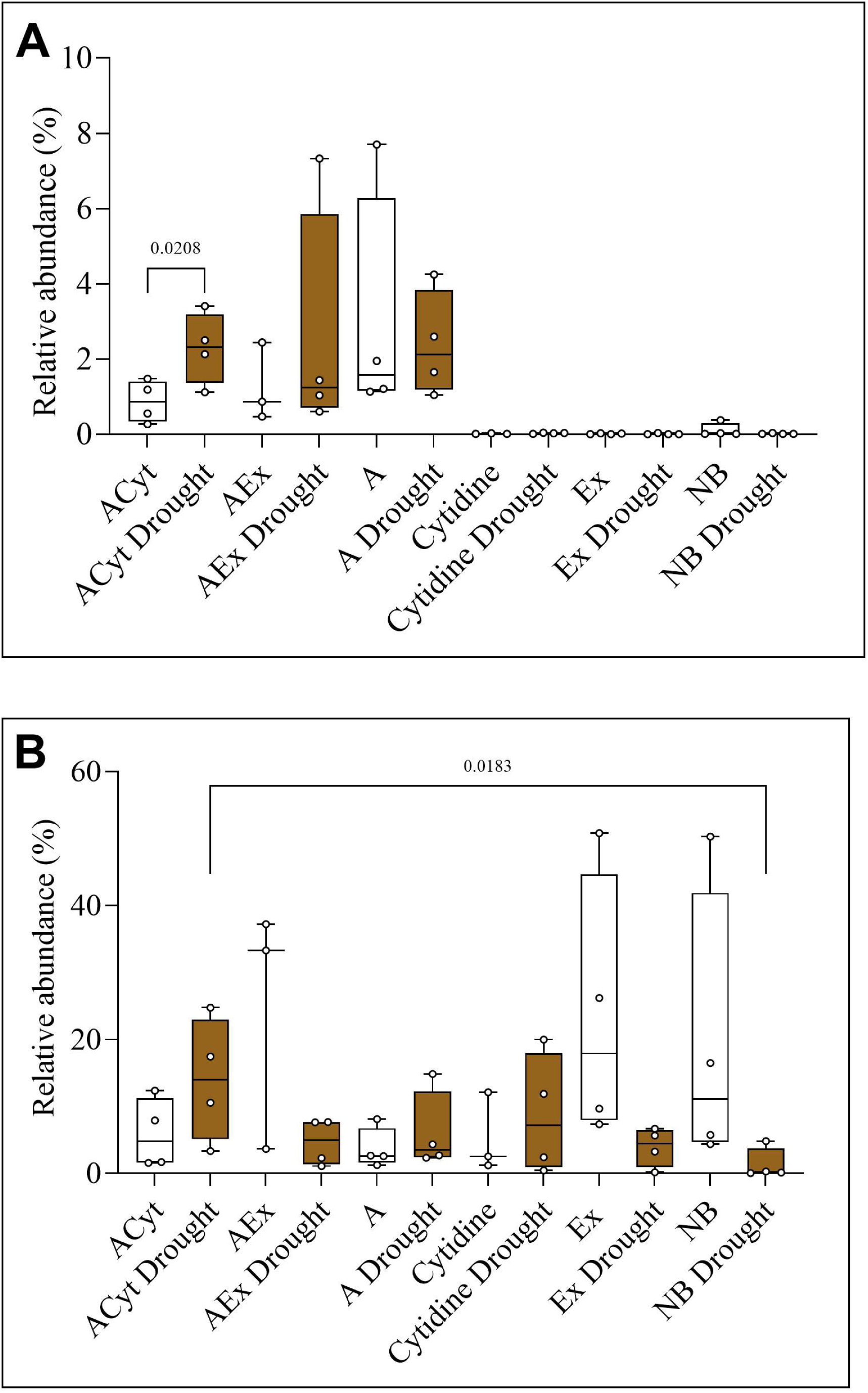
Relative abundances of *Azospirillum* 16S rDNA in the pea rhizosphere from the drought tolerance experiment. Treatment groups represent the presence or absence of *A. brasilense* Sp7 (A) and seedling exudate (Ex) or cytidine (Cyt) under watered or drought conditions. Panel A shows the relative abundance of the inoculant *A. brasilense* Sp7 (AF533354.1.1439) and Panel B shows the relative abundance of all taxa from the *Azospirillum* genus. Lines connecting two bars indicate *p*-values from pairwise comparisons with Mann-Whitney that are <0.05.

*A. brasilense* itself is a known enhancer of drought tolerance and recovery after re-watering across many crop plants (Cohen et al. 2015; Curá et al. 2017; Shelud’ko et al. 2025), some of these studies also observed an increase in shoot biomass. The effect of cytidine may be similar to the one observed for certain purines that get exuded by rice (*Oryza sativa*) under drought stress, where they are proposed to function as signaling molecules (Li et al. 2025a). The better performance of the ACyt treatment relative to the AEx one may be attributed to the much larger concentration of cytidine in the ACyt treatment relative to AEx. To summarize the phenotypic experiments, we believe that cytidine may have facilitated the establishment and/or proliferation of *A. brasilense* in the pea rhizosphere during early root colonization. Ultimately, the increased abundance of *Azospirillum* in plants from the ACyt treatment group may have amplified the published effects of *A. brasilense*.

### Alpha-diversity of the pea rhizosphere

The complete 16S rDNA data set from the pea rhizosphere was used to evaluate the impact of drought stress and treatments on alpha-diversity of the bacterial communities. Shannon index (Shannon 1948) shows a relative trend among several of the treatment groups that were inoculated with *A. brasilense* Sp7, where drought stressed groups tended to show higher diversity than their respective watered counterparts, the exception was the comparison between ACyt and ACyt Drought (Fig. 4A). This trend was not visible among any of the un-inoculated groups. Shannon index for these groups was generally somewhat lower than for inoculated groups. A similar trend was observed for Pielou (Pielou 1966) evenness (Fig. 4B). For both these indices, ACyt Drought and AEx Drought showed significantly higher values than NB Drought.

**Figure 4.**
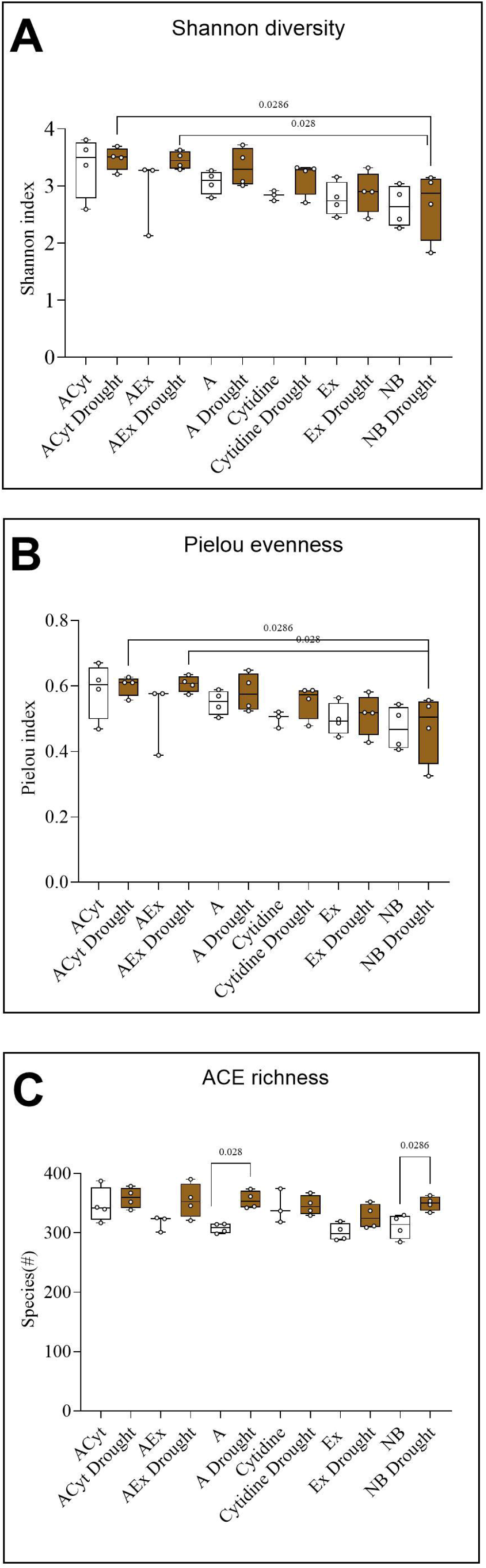
Alpha diversity of the rhizosphere bacterial community. Treatment groups represent the presence or absence of *A. brasilense* Sp7 (A) and seedling exudate (Ex) or cytidine (Cyt) under watered or drought conditions. Alpha diversity was evaluated using Shannon (A), Pielou (B), and ACE (C) indices. Lines connecting two bars indicate *p*-values from pairwise comparisons with Mann-Whitney that are <0.05.

ACE richness (Chao and Lee 1992) showed that drought stressed groups tended to show higher species richness than their respective watered counterparts (Fig. 4C). This was seen through inoculated and un-inoculated treatment groups. Two of the pairwise comparisons, A vs A Drought and NB vs NB Drought yielded *p*-values of statistical significance at percent increases in response to drought of 15.5 and 12.6%, respectively.

High biodiversity in soil is generally considered beneficial because diverse communities support a broad range of ecosystem functions, including nutrient cycling, organic matter decomposition, pathogen suppression, and resilience to environmental stress (Wagg et al. 2014; Delgado-Baquerizo et al. 2020). Compared with natural ecosystems, conventionally managed agricultural soils typically harbor less diverse and more homogeneous microbial communities, reflecting the long-term impacts of land conversion and intensive management practices (Banerjee et al. 2026). Hence, the application of PGPR has gained attention as a strategy to increase soil microbial diversity and re-establish beneficial microbial interactions in agricultural soils (Delgado-Baquerizo et al. 2020; Banerjee et al. 2026).

In this context, our data on alpha-diversity are remarkable. First, drought-stressed groups generally showed higher species richness than their watered counterparts, even though only two comparisons were statistically significant. Previous studies on the effect of drought on alpha-diversity report inconsistent effects, possibly because responses vary with host species, soil conditions, and drought severity. For example, drought increased rhizosphere bacterial richness and diversity in cowpea (*Vigna unguiculata*) (Halo et al. 2025) and alpha-diversity in cotton (*Gossypium hirsutum*) (Ullah et al. 2019). In contrast, drought had a negligible effect on rice rhizosphere alpha-diversity but clearly altered community composition (Li et al. 2023). Drought reduced rhizosphere bacterial diversity by 42% in German garlic (*Allium senescens)* (Song et al. 2025). Our own data contribute to this body of knowledge by supporting the idea that at least in some instances, drought increases species richness in soil.

Even more intriguingly, Pielou evenness increased solely by our two treatments. ACyt Drought and AEx Drought were the only two treatment groups that differed from NB Drought with statistical significance (Fig. 4B). In contrast to the phenotypic analysis, there was little difference between the two treatments, indicating that the increase in evenness can not be the sole driver behind the increase in shoot health and biomass of the drought treated plants.

### The microbiome of the pea rhizosphere

To maximize microbial functions in agroecosystems, the composition of the microbiome is important alongside alpha-diversity (Toju et al. 2018). The distribution of microbial communities across the test groups was computed from the 16S rDNA data set as percent relative abundances and presented as heatmaps. Proteobacteria were split into two sub-phyla, Alpha-and Gammaproteobacteria. These were the predominant taxonomic units, showing an almost inverse pattern of abundances across treatment groups (Fig. 5A). The drought groups ACyt Drought, AEx Drought, and A Drought had higher relative abundances of Alphaproteobacteria, when compared to the respective watered groups. Gammaproteobacteria showed the opposite pattern. Gammaproteobacteria were represented more strongly in un-inoculated groups than in inoculated ones. Notably, cytidine yielded the highest relative abundance within Gammaproteobacteria, while recording the lowest within Alphaproteobacteria.

**Figure 5.**
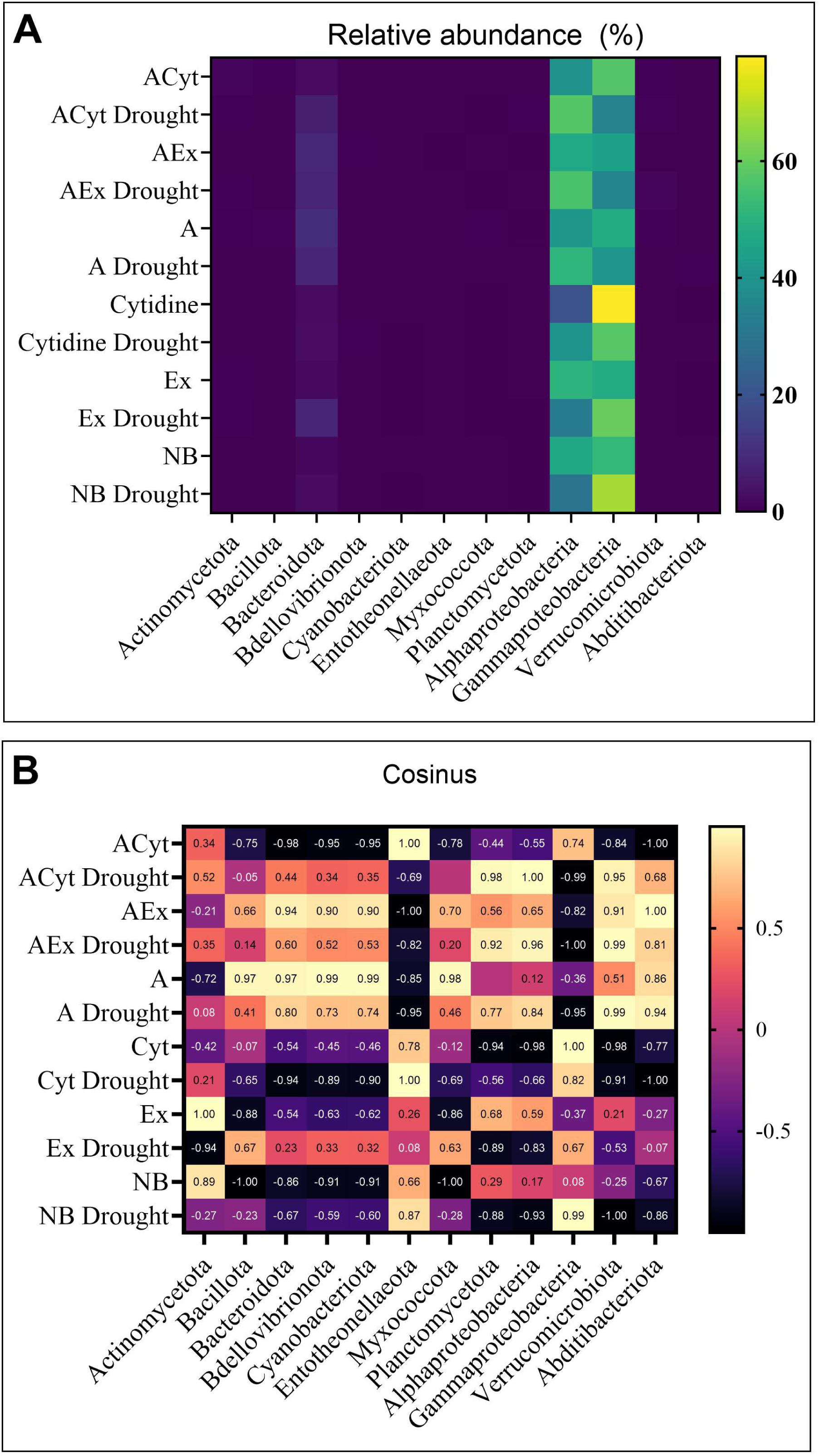
Heatmap of mean relative abundances (%) of major bacterial phyla across rhizosphere treatment groups. Each row represents a treatment group, and each column represents a bacterial phylum (or sub-phylum). In Panel A, the color gradient represents percent relative abundance. Panel B is the cosine similarity between bacterial phyla/sub-phyla and treatment groups. The color gradient indicates association strength.

To quantify association strengths between treatment groups and phyla, cosine similarity (Santos-Medellín et al. 2017) was computed from the relative abundance data (Fig. 5B). Alphaproteobacteria had strong associations with ACyt Drought, AEx Drought, and A Drought (Cos θ >0.8) and strong negative associations with Cyt, Ex Drought, and NB Drought (Cos θ <-0.8). Gamma-proteobacteria showed negative associations with ACyt Drought, AEx Drought, and A Drought and positive associations with Cyt, Cyt Drought, and NB drought. The strong abundance of Proteobacteria in our pea rhizosphere is consistent with previous reports on peas (Kichko et al. 2022), cow peas (Halo et al. 2025), and peanuts (Dai et al. 2019).

In an attempt to narrow down effected treatment taxa from phyla to genera, we separately analyzed orders within Alphaproteobacteria and Gammaproteobacteria, then families, then genera (Fig. 6). Starting with Alphaproteobacteria, we laid a path from Azospirillales and Hyphomicrobiales (left side, Fig. 6A) through Rhizobiaceae, Beijerinckiaceae, and Caulobacteraceae (left side, Fig. 6B) to *Rhizobium* and *Agrobacterium* (left side, Fig. 6C). Highest abundances of *Rhizobium* were in the ACyt Drought, AEx Drought, and A Drought groups, as well as NB Drought. This pattern of abundances across test groups was similar to that from Alphaproteobacteria. Starting with Gammaproteobacteria, there was a similar path from Enterobacterales and Burkholderiales (right side, Fig. 6A) through Erwiniaceae, Oxalobacteraceae, Burkholderaciaceae (right side, Fig. 6B) to *Pantoea* and *Massilia* (right side, Fig. 6B). The abundance pattern across treatment groups from *Massilia* was similar to that from Gammaproteobacteria. Overall, we were able to narrow down the sub-phyla of Alpha-and Gammaproteobacteria to the genera *Rhizobium* and *Massilia* as being impacted by the treatments.

**Figure 6.**
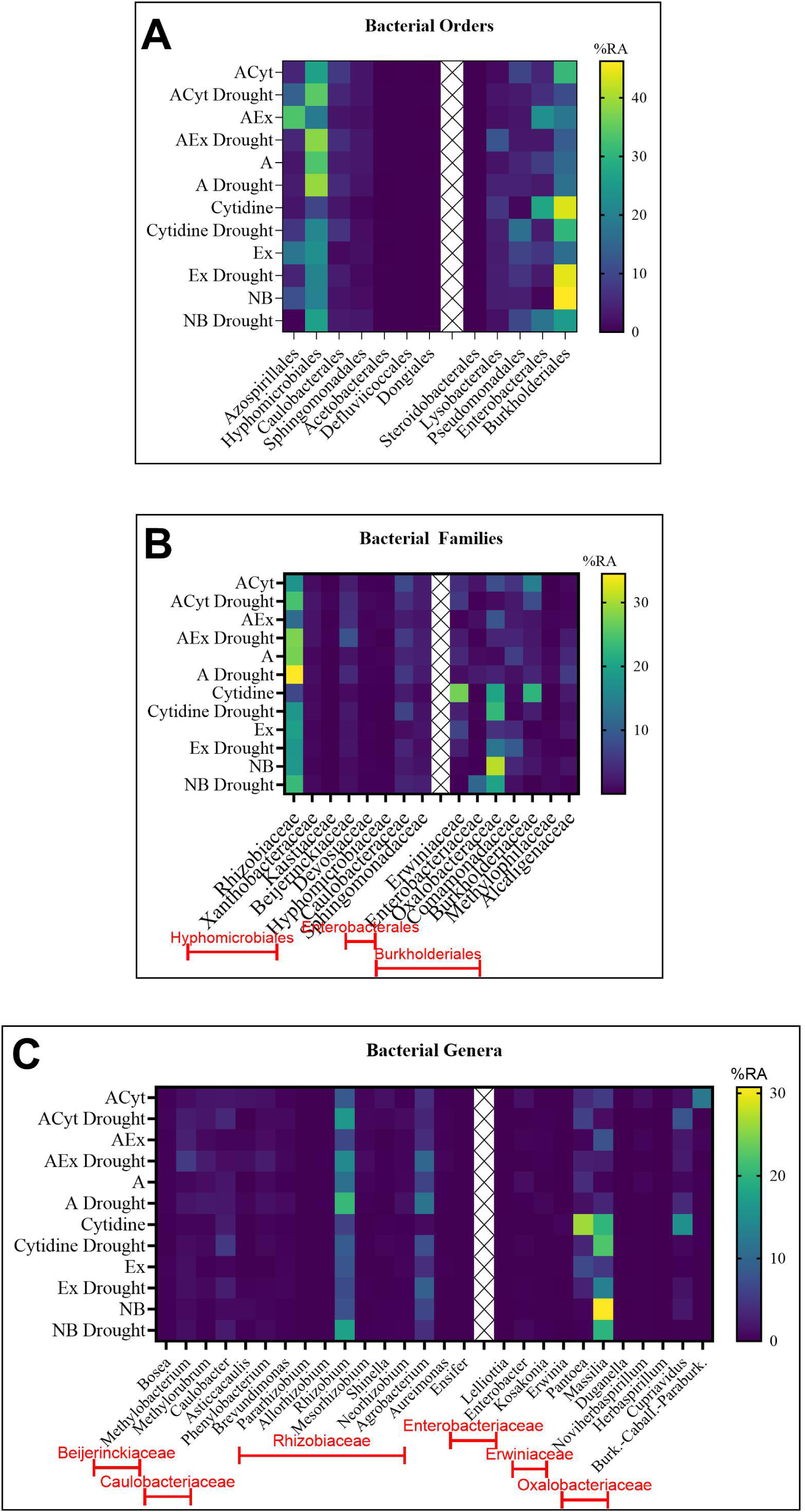
Percent relative abundances (%) of bacterial orders (A), families (B), and genera (C) across treatment groups. Each row represents a treatment group, and each column represents a bacterial order. Color intensity indicates the percentage relative abundance. The x marked areas separate Alphaproteobacteria on the left side from Gammaproteobacteria on the right side.

*Rhizobium* is a genus that was described as part of the drought response in *Medicago ruthenica* (Xing et al. 2023) and peas (Kichko et al. 2022); this, too is consistent with our study. Intriguingly, the pea study identified five rhizobial genera (*Rhizobium, Mesorhizobium, Neorhizobium, Pararhizobium*, and *Allorhizobium*) as drought responsive, as well as Xanthobacteraceae (Kichko et al. 2022). The consistency between families and genera from this study to ours is remarkable. For Xanthobacteraceae, we found that abundance of 16S rDNA was higher when *A. brasilense* Sp7 was present for all comparisons with the respective un-inoculated treatment group. Drought increased abundance especially high under the ACyt treatment (2.54% vs 1.76). For *Rhizobium*, abundance was higher under drought conditions in all three comparisons where *A. brasilense* Sp7 was present and the NB group. This was at abundances between 6% and 20%. The pattern across test groups was very similar between *Rhizobium* and Alphaproteobacteria, the latter at abundances ranging between 20% and 80%. *Pararhizobium* and *Allorhizobium* were present in our study, but at extremely low abundances (<0.03%). *Mesorhizobium* abundance was highest in the AEx Drought test group (0.96%) and very low in un-inoculated groups. *Neorhizobium* was highest in the A Drought (1.29%) and ACyt Drought (3.14%) treatment groups.

Several characteristics of *A. brasilense* Sp7 may enable the PGPR to support recruitment of native Xanthobacteraceae, *Rhizobium, Mesorhizobium,* and *Neorhizobium* to the pea rhizosphere. First, *A. brasilense* promotes root development (Cassán et al. 2009), which may provide a larger surface area for colonization by other microbes (Galindo-Castañeda et al. 2024). Second, *A. brasilense Sp7* can increased flavonoid production, which may favor compatible rhizobia by activating NodD-regulated nodulation genes (Ruan et al. 2026), thereby enhancing symbiotic interactions (Hoggard et al. 2026). Among the potential benefits of enriching the rhizosphere with *Rhizobium*, *Mesorhizobium*, and *Neorhizobium* is the ability of some of the taxa to fix nitrogen (Udvardi and Poole 2013; Lindström and Mousavi 2020). While *A. brasilense* can fix nitrogen as well, it is better recognized for all the effects that reach beyond nitrogen fixation (Fukami et al. 2018; Cassán 2026).

Like Gammaproteobacteria, *Massilia* had a higher abundance in the absence of the inoculant *A. brasilense* Sp7 and in most watered treatment groups. Note that the genus *Massilia* has recently been re-classified (Bowman 2023) and many of the species are now designated *Telluria*. *Massilia* was described as an early root colonizer in cucumbers (Ofek et al. 2012) and key driver of the drought response of *Arabidopsis thaliana* (Li et al. 2025b). As the decline of *Massilia* coincided with an increased abundance of *Rhizobium*-related taxa, we believe that we are watching a restructuring of the microbial community away from *Massilia* towards nitrogen fixing *Rhizobium*. Between *Azospirillum* and the *Rhizobium* group of rhizobacteria, the pea rhizosphere now includes bacteria that cover a wide range of physiological functions, which may make the plant ideally suited to withstand drought conditions.

## Acknowledgements

The authors thank Dr. Gladys Alexandre (University of Tennessee, Knoxville, TN) for providing us with *A. brasilense* Sp7 and Alan Wolfe (University of Loyola, Mayville, IL) and Samiran Banerjee (NDSU) for useful ideas regarding the analysis of the 16S rDNA data. Kaycie Schmidt (Dr. Glass Biotech Innovation Core) is thanked for 16S rDNA sequencing. The research was supported by a grant from the NDSU Foundation. B.M.P. was supported, in part, by the intramural research program of the USDA/NIFA, Hatch project accession number, 7009493.

## Author contributions

B.O. was responsible for data curation, formal analysis, investigation, methodology, validation, visualization, and writing-review and editing. J.A.T did investigation and writing-review and editing. S.M.H. did investigation, methology, project administration, supervision, and writing-review and editing. B.M.P. was responsible for conceptualization, funding acquisition, project administration, supervision, validation, and writing-original draft.

## Data availability

R scripts are at https:github.com/Ojha-B/Master-s-Thesis-Pea-Drought-Microbiome/tree/main. Protocols are published at dx.doi.org/10.17504/protocols.io.6nvw6pn5wgmk/v1.

## Supplementary data

Supplemental Figure 1. Shoot health scores from watered plants from the drought tolerance experiment.

Supplemental Table 1. 16S rDNA abundance table phyla

Supplemental Table 2. 16S rDNA abundance table orders

Supplemental Table 3. 16S rDNA abundance table families and genera

## Abbreviations

OTU: operational taxonomical unit;
PGPR: plant growth promoting rhizobacteria;
PCoA: principal component analysis. A, *A. brasilense* Sp7;
ACyt: *A. brasilense* Sp7 and cytidine;
AEx: *A. brasilense* Sp7 and seedling exudate.

## References

Armitage, J.P., Gallagher, A. and Johnston, A.W.B. (1988) Comparison of the chemotactic behaviour of *Rhizobium leguminosarum* with and without the nodulation plasmid. Mol Microbiol 2, 743–748.

Banerjee, S., Dasgupta, D. and van der Heijden, M.G.A. (2026) Agricultural intensification, microbial homogenization and loss of rare microbiota. Nat Rev Microbiol.

Bowman, J.P. (2023) Genome-wide and constrained ordination-based analyses of EC code data support reclassification of the species of *Massilia* La Scola et al. 2000 into *Telluria* Bowman et al. 1993, *Mokoshia* gen. nov. and *Zemynaea* gen. nov. Int J Syst Evol Microbiol 73.

Cassán, F., Maiale, S., Masciarelli, O., Vidal, A., Luna, V. and Ruiz, O. (2009) Cadaverine production by *Azospirillum brasilense* and its possible role in plant growth promotion and osmotic stress mitigation. Europ J Soil Biol 45, 12–19.

Cassán, F.D.d.B., L.; Wisniewski-Dyé, F.; Maroniche, G.A. (2026) The definitive handbook of Azospirillum: Springer.

Chao, A. and Lee, S.-M. (1992) Estimating the number of classes via sample coverage. J Amer Statist Assoc 87, 210–217.

Cohen, A.C., Bottini, R., Pontin, M., Berli, F.J., Moreno, D., Boccanlandro, H., Travaglia, C.N. and Piccoli, P.N. (2015) *Azospirillum brasilense* ameliorates the response of *Arabidopsis thaliana* to drought mainly via enhancement of ABA levels. Physiol Plant 153, 79–90.

Compton, K.K. and Scharf, B.E. (2021) Rhizobial chemoattractants, the taste and preferences of legume symbionts. Front Plant Sci 12.

Creus, C.M., Graziano, M., Casanovas, E.M., Pereyra, M.A., Simontacchi, M., Puntarulo, S., Barassi, C.A. and Lamattina, L. (2005) Nitric Oxide is involved in the *Azospirillum brasilense*-induced lateral root formation in tomato. Planta 221, 297–303.

Curá, J.A., Franz, D.R., Filosofía, J.E., Balestrasse, K.B. and Burgueño, L.E. (2017) Inoculation with *Azospirillum* sp. and *Herbaspirillum* sp. bacteria increases the tolerance of maize to drought stress. Microorganisms 5.

Dai, L., Zhang, G., Yu, Z., Ding, H., Xu, Y. and Zhang, Z. (2019) Effect of drought stress and developmental stages on microbial community structure and diversity in peanut rhizosphere soil. Int J Mol Sci 20.

Delgado-Baquerizo, M., Reich, P.B., Trivedi, C., Eldridge, D.J., Abades, S., Alfaro, F.D., Bastida, F., Berhe, A.A., Cutler, N.A., Gallardo, A., García-Velázquez, L., Hart, S.C., Hayes, P.E., He, J.Z., Hseu, Z.Y., Hu, H.W., Kirchmair, M., Neuhauser, S., Pérez, C.A., Reed, S.C., Santos, F., Sullivan, B.W., Trivedi, P., Wang, J.T., Weber-Grullon, L., Williams, M.A. and Singh, B.K. (2020) Multiple elements of soil biodiversity drive ecosystem functions across biomes. Nat Ecol Evol 4, 210–220.

Ejaz, S., Batool, S., Anjum, M.A., Naz, S., Qayyum, M.F., Naqqash, T., Shah, K.H. and Ali, S. (2020) Effects of inoculation of root-associative *Azospirillum* and *Agrobacterium* strains on growth, yield and quality of pea (*Pisum sativum L*.) grown under different nitrogen and phosphorus regimes. Scientia Horticult 270, 109401.

Fourneau, E., Pannier, M., Riah, W., Personeni, E., Morvan-Bertrand, A., Bodilis, J. and Pawlak, B. (2024) A “love match” score to compare root exudate attraction and feeding of the plant growth-promoting rhizobacteria *Bacillus subtilis*, *Pseudomonas fluorescens*, and *Azospirillum brasilense*. Frontiers in microbiology 15, 1473099.

Fukami, J., Cerezini, P. and Hungria, M. (2018) *Azospirillum*: benefits that go far beyond biological nitrogen fixation. AMB Express 8, 73.

Galindo-Castañeda, T., Hartmann, M. and Lynch, J.P. (2024) Location: root architecture structures rhizosphere microbial associations. J Experim Botany 75, 594–604.

Gaworzewska, E.T. and Carlile, M.J. (1982) Positive chemotaxis of *Rhizobium leguminosarum* and other bacteria towards root exudates from legumes and other plants. Microbiology 128, 1179–1188.

Haghpanah, M., Hashemipetroudi, S., Arzani, A. and Araniti, F. (2024) Drought tolerance in plants: physiological and molecular responses. Plants 13, 2962.

Halo, B.A., Aljabri, Y.A.S. and Yaish, M.W. (2025) Drought-induced microbial dynamics in cowpea rhizosphere: Exploring bacterial diversity and bioinoculant prospects. PLoS One 20, e0320197.

Hoggard, S., Calhoun, M., Bommes, A., Rahmatallah, Y., Neuhaus, D., Glazko, G. and Mukherjee, A. (2026) Diffusible signals from *Azospirillum brasilense* promote rice growth and induce distinct gene expression changes in plant roots. Front Plant Sci Volume 17 - 2026.

Kichko, A.A., Gladkov, G.V., Ulianich, P.S., Safronova, V.I., Pinaev, A.G., Sekste, E.A., Belimov, A.A. and Andronov, E.E. (2022) Water stress, cadmium, and plant genotype modulate the rhizosphere microbiome of *Pisum sativum* L. Plants (Basel) 11.

Li, G., Wang, K., Qin, Q., Li, Q., Mo, F., Nangia, V. and Liu, Y. (2023) Integrated microbiome and metabolomic analysis reveal responses of rhizosphere bacterial communities and root exudate composition to drought and genotype in rice (*Oryza sativa* L.). Rice (N Y) 16, 19.

Li, Y., Ye, Z., Xiang, J., Li, S., Zheng, Z., Li, Y., Fang, Y., Zhang, X., Chen, X. and Xue, D. (2025a) Purine nucleotide metabolism response to drought stress in rice. Plant Growth Regul 105, 821–832.

Li, Z., Wang, Z., Zhang, Y., Yang, J., Guan, K. and Song, Y. (2025b) Identification of stress-alleviating strains from the core drought-responsive microbiome of *Arabidopsis* ecotypes. ISME J 19.

Lindström, K. and Mousavi, S.A. (2020) Effectiveness of nitrogen fixation in rhizobia. Microb Biotechnol 13, 1314–1335.

Nikodemova, M., Holzhausen, E.A., Deblois, C.L., Barnet, J.H., Peppard, P.E., Suen, G. and Malecki, K.M. (2023) The effect of low-abundance OTU filtering methods on the reliability and variability of microbial composition assessed by 16S rRNA amplicon sequencing. Front Cell Infect Microbiol 13, 1165295.

Nisha, F.A., Horne, S.M. and Prüß, B.M. (2025) *Azospirillum brasilense* and cytidine enhance lateral roots of peas. FEMS Microbiol Lett 372.

Nisha, F.A., Tagoe, J.N.A., Pease, A.B., Horne, S.M., Ugrinov, A., Geddes, B.A. and Prüß, B.M. (2024) Plant seedlings of peas, tomatoes, and cucumbers exude compounds that are needed for growth and chemoattraction of *Rhizobium leguminosarum* bv. *viciae* 3841 and *Azospirillum brasilense* Sp7. Can J Microbiol 70, 150–162.

O’Neal, L., Vo, L. and Alexandre, G. (2020) Specific root exudate compounds sensed by dedicated chemoreceptors shape *Azospirillum brasilense* chemotaxis in the rhizosphere. Appl Environm Microbiol 86, e01026–01020.

Ofek, M., Hadar, Y. and Minz, D. (2012) Ecology of root colonizing *Massilia* (Oxalobacteraceae). PloS one 7, e40117.

Pielou, E.C. (1966) The measurement of diversity in different types of biological collections. J Theoret Biol 13, 131–144.

Ruan, Y., Dong, S., Jiang, S., Wang, Y., Wu, X., Zhuang, Y., Wu, W., East, A.K., Xu, P., Poole, P.S., Zhang, Y. and Murray, J.D. (2026) The molecular basis of the binding and specific activation of rhizobial NodD by flavonoids. Science 391, 184–189.

Santos-Medellín, C., Edwards, J., Liechty, Z., Nguyen, B. and Sundaresan, V. (2017) Drought stress results in a compartment-specific restructuring of the rice root-associated microbiomes. mBio 8.

Shannon, C.E. (1948) A mathematical theory of communication. Bell System Techn J 27, 379–423.

Shelud’ko, A., Volokhina, I., Mokeev, D., Telesheva, E., Filip’echeva, Y., Burov, A., Borisov, I., Shirokov, A., Matora, L. and Petrova, L. (2025) Multilevel analysis of *Azospirillum* biofilms and cystlike forms and characterization of their possible role in wheat drought tolerance. Rhizosphere 33, 101029.

Song, X., Li, H., Fu, C., Li, J., Xiang, J., Sun, X., Liu, J. and Qin, L. (2025) Diversity changes of rhizosphere and endophytic bacteria in Allium senescens L. under drought stress and rewatering. Front Plant Sci 16, 1571736.

Sorensen, J.N., Edelenbos, M. and Wienberg, L. (2002) Drought effects on green pea texture and related physical-chemical properties at comparable maturity. J Amer Soc Horticult Sci 128, 128–135.

Tagoe, J., Ojha, B., Horne, S. and Prüß, B. (2026) *Azospirillum brasilense* and tomato exudate or cytidine increase phytopathogen resistance and modulate phyllosphere/rhizosphere. Can J Microbiol 72, 1–14.

Tarrand, J.J., Krieg, N.R. and Döbereiner, J. (1978) A taxonomic study of the *Spirillum lipoferum* group, with descriptions of a new genus, *Azospirillum* gen. nov. and two species, Azospirillum lipoferum (Beijerinck) comb. nov. and Azospirillum brasilense sp. nov. Can J Microbiol 24, 967–980.

Toju, H., Peay, K.G., Yamamichi, M., Narisawa, K., Hiruma, K., Naito, K., Fukuda, S., Ushio, M., Nakaoka, S., Onoda, Y., Yoshida, K., Schlaeppi, K., Bai, Y., Sugiura, R., Ichihashi, Y., Minamisawa, K. and Kiers, E.T. (2018) Core microbiomes for sustainable agroecosystems. Nature Plants 4, 247–257.

Udvardi, M. and Poole, P.S. (2013) Transport and metabolism in legume-rhizobia symbioses. Annu Rev Plant Biol 64, 781–805.

Ullah, A., Akbar, A., Luo, Q., Khan, A.H., Manghwar, H., Shaban, M. and Yang, X. (2019) Microbiome diversity in cotton rhizosphere under normal and drought Conditions. Microb Ecol 77, 429–439.

Wagg, C., Bender, S.F., Widmer, F. and van der Heijden, M.G. (2014) Soil biodiversity and soil community composition determine ecosystem multifunctionality. Proc Natl Acad Sci U S A 111, 5266–5270.

Wang, J. and Xu, Z. (2026) Plant response to and recovery from drought. Current Biology 36, R343–R362.

Wang, T., Singh, B.K., Trivedi, P., Islam, Z.F., He, J.-Z. and Hu, H.-W. (2026) Bacterial motility in rhizosphere colonization: mechanisms, constraints, and implications for microbial inoculants. ISME J 20.

Warkentin, T.V.A.B., S.; Slinkard, A. (2004) CDC Striker field pea. Can J Plant Sci, 239–240.

Xing, J., Fan, W., Wang, J. and Shi, F. (2023) Variety-driven effect of rhizosphere microbial-specific recruitment on drought tolerance of *Medicago ruthenica* (L.). Microorganisms 11.

Zheng, C., Bochmann, H., Liu, Z., Kant, J., Schrey, S.D., Wojciechowski, T. and Postma, J.A. (2023) Plant root plasticity during drought and recovery: What do we know and where to go? Front Plant Sci 14, 1084355.

