## Supplementary figures and images for "*Azospirillum brasilense* and cytidine as combination treatment to enhance drought resilience of peas"

### Fig. S1

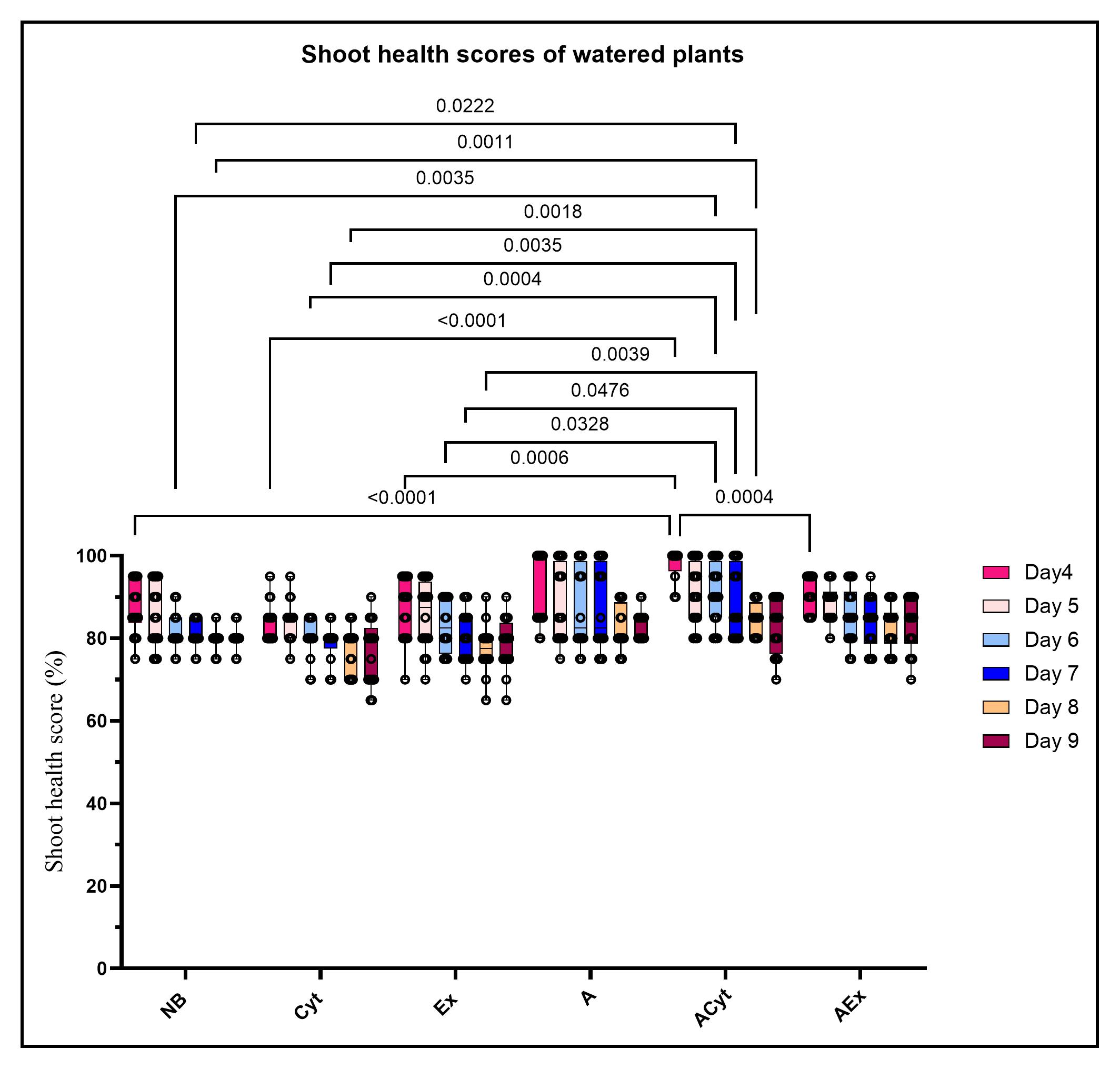
